# The plakin domain of *C. elegans* VAB-10/plectin acts as a hub in a mechanotransduction pathway to promote morphogenesis

**DOI:** 10.1101/681072

**Authors:** Shashi Kumar Suman, Csaba Daday, Teresa Ferraro, Thanh Vuong-Brender, Saurabh Tak, Sophie Quintin, François Robin, Frauke Gräter, Michel Labouesse

## Abstract

Mechanical forces control many cellular processes by eliciting a mechanotransduction response in target cells. The initial steps of mechanotransduction at hemidesmosomes remain undefined in contrast to focal adhesions and adherens junctions. Here, we focus on the *C. elegans* plectin homolog VAB-10A, the only evolutionary conserved hemidesmosome component. In *C. elegans*, muscle contractions induce a mechanotransduction pathway in the epidermis through hemidesmosomes. We used CRISPR to precisely remove spectrin repeats (SR) or a partially hidden Src-homology-3 (SH3) domain within the VAB-10 plakin domain. Deleting the SH3 or SR8 domains in combination with mutations affecting mechanotransduction, or just part of SR5 shielding the SH3 domain induced embryonic elongation arrest because hemidesmosomes collapse. Notably, recruitment of GIT-1, the first mechanotransduction player, requires the SR5 domain and the hemidesmosome transmembrane receptor LET-805. Furthermore, Molecular Dynamics simulations confirmed that forces acting on VAB-10 can render the central SH3 domain, otherwise in contact with SR4, available for interaction. Collectively, our data strongly argue that the plakin domain plays a central role in mechanotransduction and raise the possibility that VAB-10/plectin might act as a mechanosensor.

**Summary statement:** CRISPR-derived deletions reveal the roles of three spectrin repeats and an atypical SH3 domain from the plakin domain of the VAB-10 hemidesmosome component in mechanotransduction during *C. elegans* morphogenesis

## Introduction

Cells are constantly exposed to various mechanical forces, and their ability to respond is critical for tissue homeostasis, particularly in the context of morphogenesis and cancer progression. For instance during tissue and organ formation, epithelial tissues get strongly deformed due to both cell-intrinsic and cell-extrinsic forces (Gilmour et al., 2017). Yet, cells within the epithelium must maintain cell-cell and cell-matrix cohesion. The adaptation process to these forces relies on mechanosensing and the transduction of specific signals from junctions to the cytoskeleton or to the nucleus (Iskratsch et al., 2014; Ladoux et al., 2015).

Focal adhesions and adherens junctions play a central role in relaying mechanical forces. Tension exerted on those junctions result in similar effects, whereby a protein acting as a junction-linked mechanosensor gets unfolded and recruits additional proteins that either strengthen the junction or induce biochemical signalling (Chen et al., 2017; Moore et al., 2010). Typically, tension applied to focal adhesions can unfold talin, which then recruits vinculin (del Rio et al., 2009), or the adaptor p130Cas, which subsequently activates the small GTPase Rap1 (Sawada and Sheetz, 2002; Sawada et al., 2006). Likewise, α-catenin acts as an adherens junction mechanosensor that also recruits vinculin to strengthen the junction (le Duc et al., 2010; Yao et al., 2014; Yonemura et al., 2010).

Hemidesmosomes represent another mechano-sensitive junction about which much less is known. In *C. elegans*, we previously found that the hemidesmosome-like junction present in epidermal cells transmits mechanical tension exerted by muscles when they contract (Zhang et al., 2011). *C. elegans* muscles are positioned along the anterior-posterior axis under the dorso-ventral epidermis (see cross-hatched red lines in Fig. 1A-A’). They are tightly connected to the epidermis through a complex structure acting as trans-epithelial tendons, called fibrous organelles, consisting in two hemidesmosome-like junctions (CeHDs) connected by intermediate filaments (Fig. 1B) (Francis and Waterston, 1991; Gieseler et al., 2017; Vuong-Brender et al., 2016). These structures share the plectin homologue VAB-10A and intermediate filaments with the canonical vertebrate hemidesmosomes (Zhang and Labouesse, 2010). Due to this tight connection, when muscles contract, they submit dorsal and ventral epidermal cells to mechanical tension. In turn, it initiates a mechanotransduction process in the dorsal and ventral epidermal cells requiring functional hemidesmosomes (Fig. 1A-B) (Zhang et al., 2011). We previously found that the first detectable step in this mechanotransduction process corresponds to the recruitment of the adaptor protein GIT-1 to hemidesmosomes (Fig. 1C) (Zhang et al., 2011). Although VAB-10A/plectin is essential for hemidesmosome integrity (Bosher et al., 2003), its potential role in the mechanotransduction process had not been investigated. In vertebrates, plectin, a core component of vertebrate hemidesmosomes, interacts with dystroglycan to mediate mechanosignaling in lung epithelial cells, independently of hemidesmosomes, in a process leading to ERK1/2 and AMPK activation (Takawira et al., 2011). Plectin is also important to maintain the nuclear shape of keratinocytes as they adhere and spread out on micropatterns, which they do by regulating the density of the perinuclear keratin meshwork and in part through a negative regulation of MAPK activity; as such it regulates nuclear mechanotransduction (Almeida et al., 2015).

**Figure 1.**
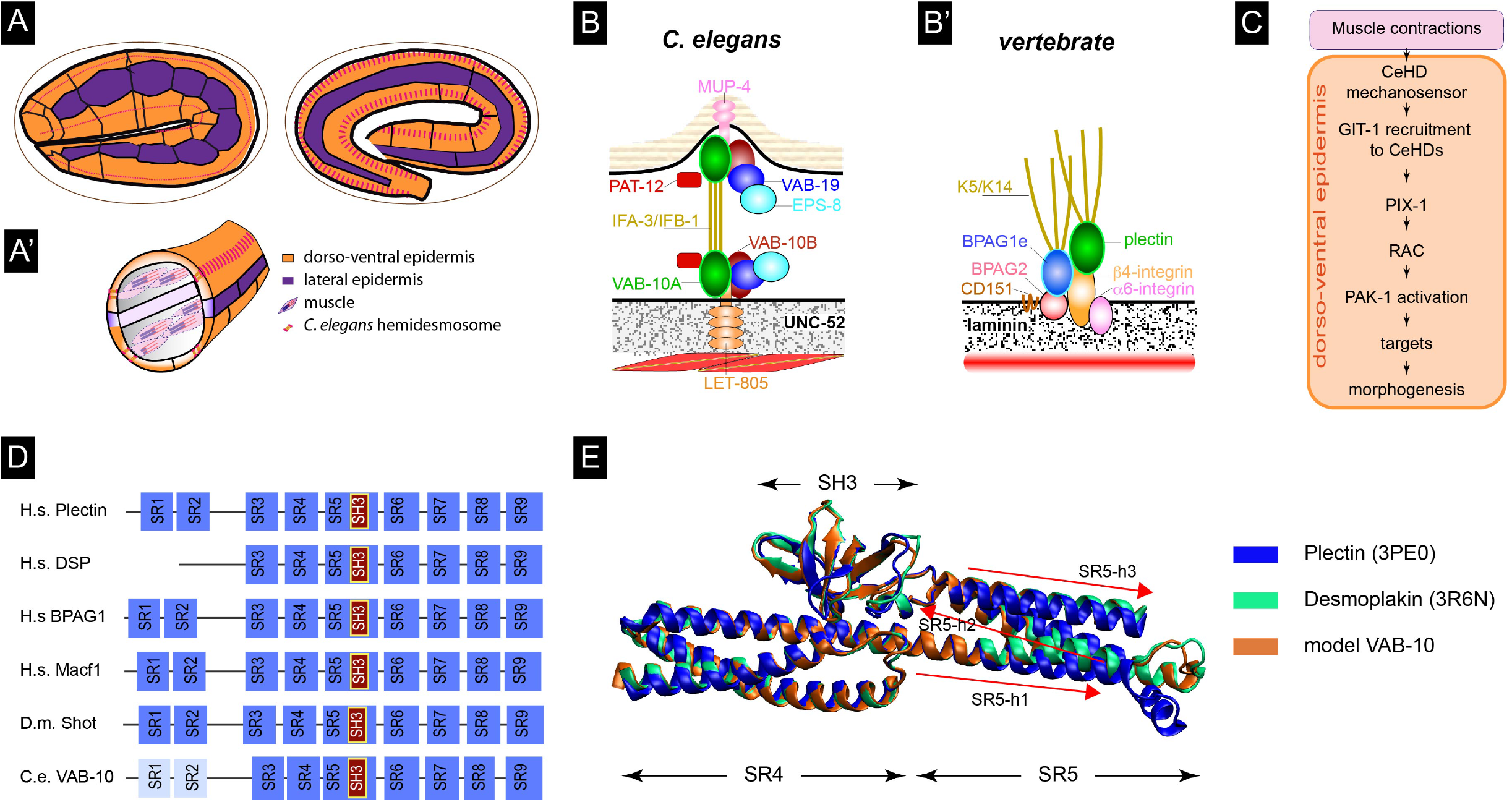
The SR4-SR5-SH3 domains of plectin and VAB-10 have a similar conformation. **(A)** Muscles are required for the embryo to elongate from the 2-fold stage (left) to the terminal 4-fold/prezel stage (right). **(A’)** Cross-section of the embryo showing the relative positions of muscles and epidermal cells. **(B)** Comparison of the embryonic *C. elegans* and **(B’)** vertebrate hemidesmosome junctions. The drawing is only showing essential *C. elegans* hemidesmosome proteins (GIT-1, PIX-1, PAK-1 not included). Besides intermediate filaments, the only conserved protein in common is the plakin: Plectin and BPAG1e in vertebrates, VAB-10A in *C. elegans* (green; the VAB-10B/MACF plakin transiently associates with the hemidesmosome). LET-805 basally, MUP-4 apically, the intermediate filament dimers IFA-3/ IFB-1, PAT-12 are nematodespecific proteins; VAB-19 and its binding partner EPS-8B have homologs in vertebrates (Kank1 and EPS8, respectively) which are not known to associate with vertebrate hemidesmosomes; the basal ECM ligand UNC-52 is homologous to perlecan, whereas the apical one includes several proteins with Zona Pellucida domains. **(C)** Outline of the mechanotransduction pathway occurring in the epidermis in response to the tension created by the contraction of muscles, which are tightly linked to the epidermis. **(D)** Comparison of the plakin domain of plectin and spectraplakins, showing that all of them have a predicted SH3 domain nested within a spectrin domain. The structure prediction is based on (Choi and Weis, 2011; Daday et al., 2017; Ortega et al., 2016), and BLASTP alignments. Numbering of the SR domains is based on plectin in this and all subsequent figures (Ortega et al., 2011; Ortega et al., 2016). **(E)** 3D structure the SH3 and surrounding spectrin repeats from human Plectin (PDB 3PE0; blue), human desmoplakin (PDB 3RN6; green) and superimposed Pymol prediction for the homologous spectrin repeat and SH3 domain from VAB-10 (orange). The only major difference between them is in the minor helix on the right.

The goal of the present work was first to examine whether VAB-10A was required to mediate the mechanotransduction response triggered in epidermal cells by the periodic contraction of muscles. We sought to determine the domains that could mediate this response, by removing specific domains within VAB-10, using a CRISPR/Cas9 approach. Our data reveal a key role for an SH3 protein-protein interaction domain and its spectrin repeat shield, within the plakin domain, in mediating mechanotransduction. Since VAB-10A/plectin is the only hemidesmosomal protein conserved between *C. elegans* and vertebrates (Zhang and Labouesse, 2010), it is likely that responses gained from *C. elegans* will apply to vertebrate hemidesmosomes.

## Results

### Isolation of novel SH3 and spectrin repeat mutations in *vab-10*

Proteins found at *C. elegans* hemidesmosomes are detailed in Fig. 1B. As outlined above, VAB-10A is the only conserved hemidesmosome protein between *C. elegans* and vertebrates (Zhang and Labouesse, 2010) - note that the gene *vab-10* produces two major isoforms through alternative splicing, VAB-10A/plectin and VAB-10B/MACF, which have the N-terminal actin-binding domain and the plakin domain in common (see below) (Bosher et al., 2003). Several considerations prompted us to more specifically focus on the potential role of the central plakin domain. First, this domain is well conserved among plakin and spectraplakin families such as vertebrate desmoplakin, plectin and MACF/ACF7, *Drosophila* Shot and *C. elegans* VAB-10, containing up to nine spectrin repeats (SR) (Zhang et al., 2017). Interestingly these domains are known to unfold *in vitro* under mechanical stress (Law et al., 2003; Lenne et al., 2000). Second, the central SR harbors a predicted SH3 domain (Fig. 1D), although it lacks some critical residues compared to canonical SH3 domains questioning its ability to interact with Pro-rich regions (Choi and Weis, 2011; Ortega et al., 2011). Indeed, the crystal structure of that region from desmoplakin and plectin suggests that the SH3 domain interacts with the upstream spectrin repeat to stabilize the plakin domain, which should make the SH3 binding region partially occluded and prevented from interacting with other proteins (Fig. 1E) (Choi and Weis, 2011; Ortega et al., 2011). Third, recent molecular dynamics simulations (MDS) on plectin and desmoplakin suggested that mechanical force should unfold the two neighboring spectrin repeats to unmask the SH3 domain (Daday et al., 2017).

We first modeled the equivalent region of VAB-10 on top of human plectin, which showed that its SH3 domain should also be involved in making contacts with the upstream SR4 repeat (Fig. 1E). To functionally test the importance of the plakin domain, we engineered several mutations in *vab-10* by CRISPR-mediated recombination (Fig. 2A-B). Specifically, we introduced a deletion of the first two alpha-helices within the SR5 preceding the SH3 domain, named *vab-10*(*mc64*) or *vab-10*(Δ*SR5h1-h2*), a deletion of the entire SH3 domain named *vab-10*(*mc62*) or *vab-10*(Δ*SH3*), deletions removing SR7 or SR8 - named *vab-10*(*mc97*) and *vab-10*(*mc98*), respectively (or Δ*SR7* and Δ*SR8*) - deletion of the SH3 PSVV residues contributing to ligand binding among canonical SH3 domains, named *vab-10*(*mc56*), and several point mutations in conserved SH3 residues or in a cysteine of SR4 predicted to interact with SH3 (Fig. 2A-B, Fig. S1). Our attempts to isolate similar deletions of the other SR domains were unfortunately unsuccessful. By comparison with previously known *vab-10* alleles, *vab-10*(Δ*SR5h1-h2*) behaved like a strong *vab-10* allele, segregating 100% dead embryos that elongated slightly beyond the 2-fold stage (Fig. 2C-D, Table 1). By contrast, over 90% animals homozygous for *vab-10*(Δ*SH3*), deletion of the SH3 PSVV residues, *vab-10*(Δ*SR7*), *vab-10*(Δ*SR8*) were normal, with a few late embryo/arrested larvae displaying a very lumpy morphology (Fig. 2E-G). Other point mutations were homozygous viable with no apparent phenotype (Fig. 2I; Table S1). In the remainder of this study, we will mainly focus on the SH3 deletion, SR8 deletion and on the deletion removing the two SR5 alpha-helices preceding the SH3 domain, and briefly mention the results obtained with other mutants.

**Figure 2.**
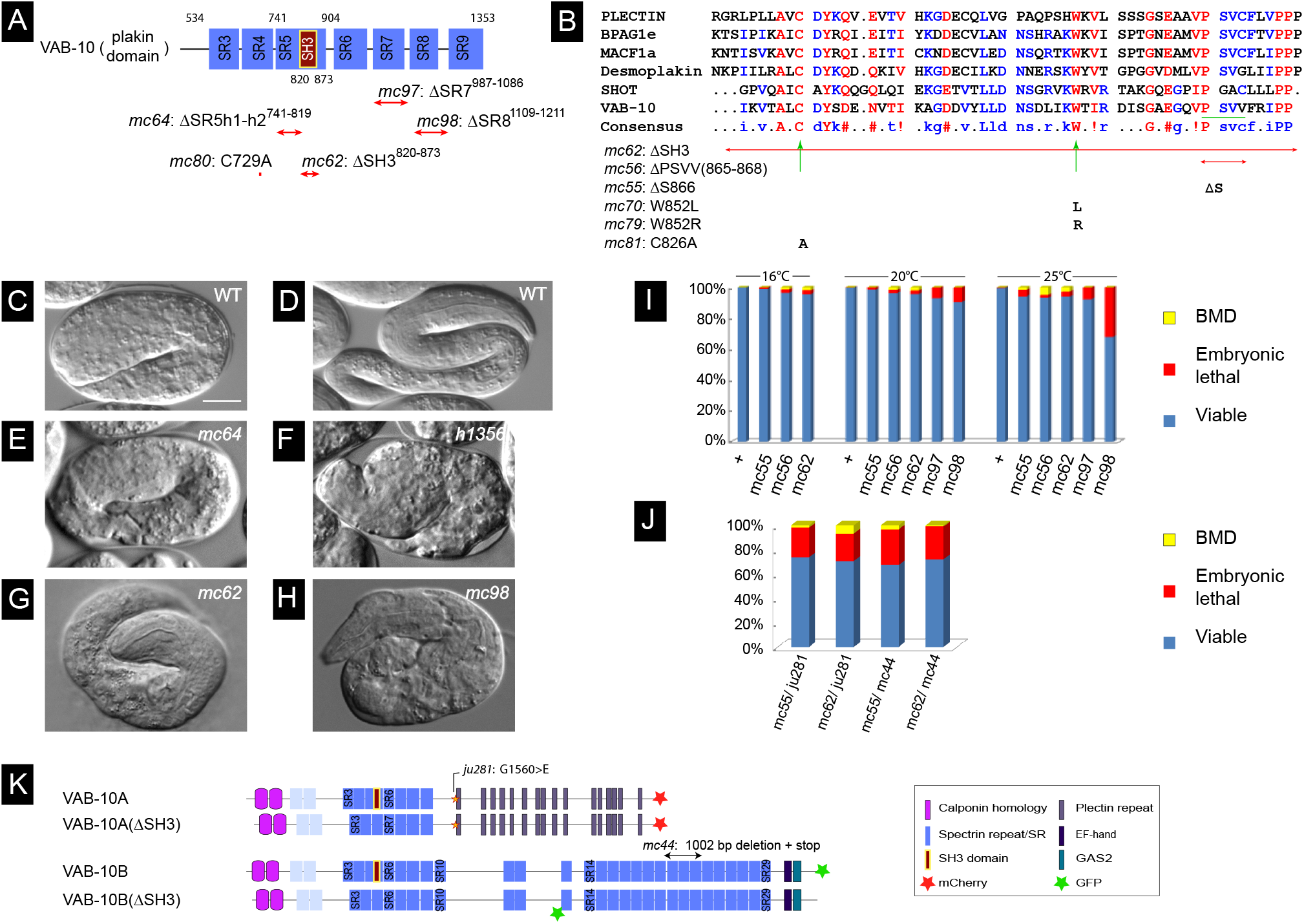
The SR5 and SH3 domains of VAB-10 are important for embryonic elongation. **(A)** Position of the CRISPR-generated mutations in the plakin domain. **(B)** Alignment of the SH3 domain among spectraplakins and plakins (as predicted by the SMART software), along with the positions of the residues mutated by CRISPR. **(C-H)** DIC micrographs of a wild-type 2-fold embryo (**C**) and young L1 hatchling (**D**), an arrested *mc64* (noted Δ*SR5h1-h2* in the text) embryo (**E**), a *h1356* embryo which is a presumptive *vab-10* null allele (**F**), a rare malformed *mc62* embryo (noted Δ*SH3* in the text) (**G**) and a rare malformed *mc98* embryo (noted Δ*SR8* in the text) (**H**). **(I)** Quantification of the embryonic and larval lethality rate among the main new *vab-10* alleles (see also Table 1). **(J)** Allelic complementation tests between *mc55* or *mc62* and the very strong *vab-10A*(*ju281*) and *vab-10B*(*mc44*) alleles; the phenotypes correspond to those observed in the progeny of *mc62/ju281* and *mc62/mc44* trans-heterozygous adults. Chi-square analysis shows the distributions are not statistically different. **(K)** The VAB-10A and VAB-10B isoforms were tagged with mCherry and GFP, respectively. Additionally, these markers were introduced in a *vab-10*(Δ*SH3*) mutant, except that the GFP could not be obtained at the very C-term for *vab-10B*(Δ*SH3*). Scale bars, 10 μm.

**Table 1.**
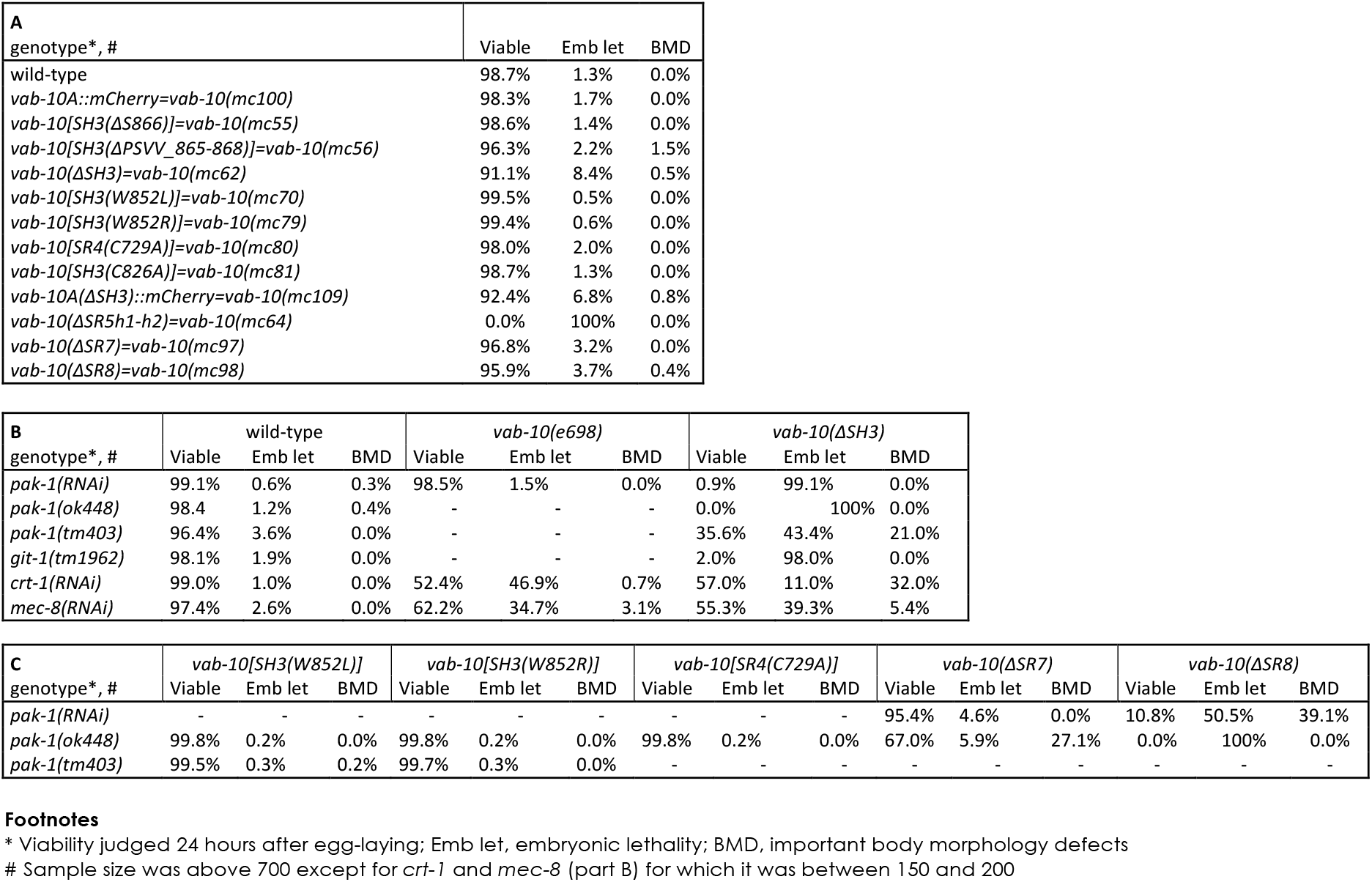
Phenotypes of mutations affecting the VAB-10 plakin domain

To test whether *vab-10*(Δ*SH3*) affects the *vab-10A* or *vab-10B* isoforms, we performed complementation tests with the embryonic lethal alleles *vab-10A*(*ju281*) and *vab-10B*(*mc44*) (see Fig. 2I for their positions). The prediction if *vab-10*(Δ*SH3*) disrupts a specific *vab-10* isoform is that the corresponding trans-heterozygous combination should not be viable or should display serious larval defects, whereas the other trans-heterozygous combination should be normal. In contrast to what we had previously observed when testing the complementation between *vab-10A*(*e698*) and other *vab-10* alleles (Bosher et al., 2003), we could establish trans-heterozygous *vab-10A*(*ju281*)/*vab-10*(Δ*SH3*) and *vab-10B*(*mc44*)/*vab-10*(Δ*SH3*) animals that both segregated between 23% and 29% embryonic lethality and up to 7% strong larval body morphology defects (Fig. 2J). Since homozygous *vab-10A*(*ju281*) or *vab-10B*(*mc44*) represent 25% of the progeny, and homozygous *vab-10*(Δ*SH3*) another 25% of the progeny of which less than 10% display strong defects (see Table1, Fig. 2I), we conclude that *vab-10*(Δ*SH3*) does not severely compromise VAB-10 function. Since the distribution of the different categories was not statistically different in the progeny of *ju281*/*mc62* versus *mc44*/*mc62* trans-heterozygotes, this complementation test alone does not reveal whether *vab-10*(Δ*SH3*) is a *vab-10A* or *vab-10B* allele, or possibly whether both isoforms might fulfill the function provided by the SH3 domain during embryonic elongation.

To facilitate the study of VAB-10, we also generated a CRISPR knock-in of the VAB-10A isoform marked by mCherry at its C-terminus as well as a ΔSH3 version of this knock-in; likewise, we generated a CRISPR knock-in of the other major isoform VAB-10B marked by GFP at the C-ter for the wild-type form, and internally before its SR13 for the ΔSH3 form (Fig. 2I). Homozygous VAB-10A::mCherry and VAB-10B::GFP animals were normal, whereas their ΔSH3 derivatives behaved like *vab-10*(Δ*SH3*) animals, segregating <10% arrested embryos or L1 larvae (Table 1).

### The VAB-10A plakin domain is essential for mechanotransduction

We had previously found that mutations in *git-1* or *pak-1* induce high levels of embryonic lethality when combined with the weak allele *vab-10A*(*e698*), which affects a non-conserved region of the protein, resulting in compromised hemidesmosomes (Zhang et al., 2011).

We thus tested whether double mutants between *vab-10*(Δ*SH3*) or *vab-10*(Δ*SR8*) and *git-1*(*tm1962*) or *pak-1*(*ok448*) would induce a similar phenotype. Strikingly, time-lapse DIC microscopy of *vab-10*(Δ*SH3*); *pak-1*(*RNAi*), *vab-10*(Δ*SH3*); *git-1*(*tm1962*) or *vab-10*(Δ*SR8*); *pak-1*(*RNAi*) embryos showed that most of them did not progress beyond the 2.5-fold stage (Fig. 3A-C; Movies S1-S3). Specifically, 98% *vab-10*(Δ*SH3*); *git-1*(*tm1962*), 100% *vab-10*(Δ*SH3*); *pak-1*(*ok448*) and *vab-10*(Δ*SR8*); *pak-1*(*ok448*) embryos failed to hatch (Table 1, Fig. 3B), underlining a strong synergistic effect between mutations that have on their own very weak embryonic defects. The allele *pak-1*(*ok448*) is a presumptive null removing the kinase domain; note that inducing *pak-1* knockdown by RNAi in the *vab-10*(Δ*SR8*) background induced 50% embryonic and 40% larval lethality which is not as penetrant as with the *pak-1*(*ok448*) allele (Table 1). These double mutants elongated at a wild-type rate until the 2.2-fold stage, then arrested (Fig. 3A,C). A double mutant combining *vab-10*(Δ*SH3*) with a mutant removing the GTPase-binding domain of PAK-1, *pak-1*(*tm403*), was not as severely affected (Table 1). We also observed approximately 50% lethality after RNAi against *mec-8* or *crt-1* in the *vab-10*(Δ*SH3*) background, two strong enhancers of *vab-10A*(*e698*) without an embryonic phenotype of their own (Zahreddine et al., 2010) (Table 1). These genetic interactions, together with the type of embryonic arrest, strongly support the notion that *vab-10*(Δ*SH3*) and *vab-10*(Δ*SR8*) behave like *vab-10A* alleles. By contrast, *vab-10*(Δ*SR7*) induced less significant lethality when combined with *pak-1*(*ok448*) (Fig. S1; Table 1). Hence, based on genetic interactions with genes known to mediate mechanotransduction, the SH3 and SR8 domains, hence a large part of the plakin domain are important for mechanotransduction.

**Figure 3.**
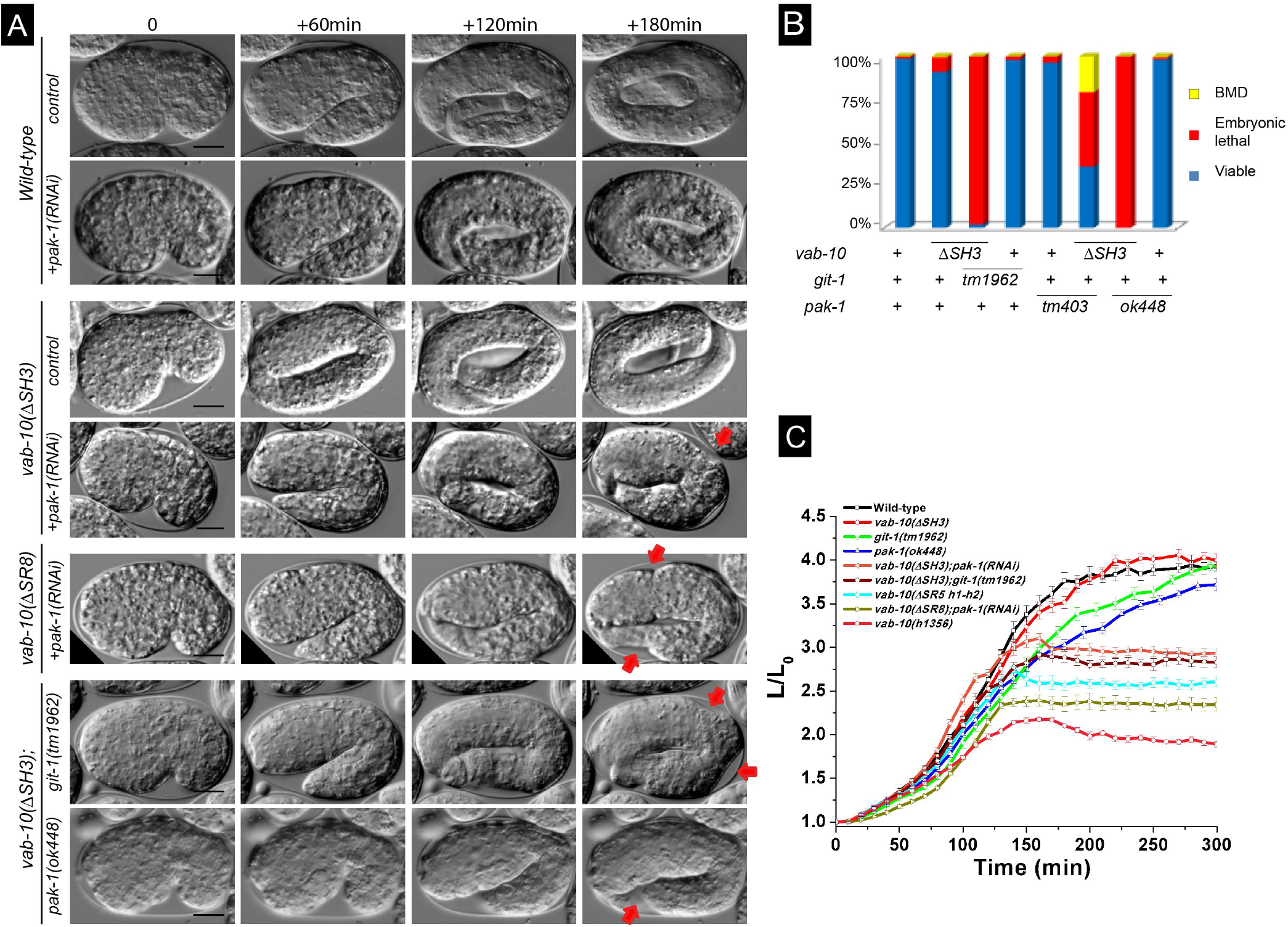
Strong synergistic interactions between novel *vab-10* alleles and mutations affecting the mechanotransduction pathway. **(A)** DIC pictures showing the elongation of control embryos (row 1), of *pak-1*(*RNAi*) embryos (row 2), *vab-10A*(Δ*SH3*) embryos (row 3), and *vab-10A*(Δ*SH3*) embryos after *pak-1*(*RNAi*) (row 4), *vab-10A*(Δ*SR8*); *pak-1*(*RNAi*) embryos (row 5), *vab-10A*(Δ*SH3*); *git-1*(*tm1962*) embryos (row 6), *vab-10A*(Δ*SH3*); *pak-1*(*ok448*) embryos (row 7); pictures from rows 2, 4 and 6 are Movies S1-S2-S3, respectively. The time interval between images is indicated at the top. Red arrow, localized irregularity in the body wall. **(B)** Quantification of the embryonic and larval lethality in the double mutants shown in (**A**). **(C)** Elongation curves of the same double mutants. Scale bars, 10 μm.

### The VAB-10A plakin domain contributes to maintain hemidesmosome integrity

We had previously found that hemidesmosomes fall apart in most *vab-10A*(*e698*); *pak-1*(*ok448*) double-mutants (Zhang et al., 2011). To define whether embryos having plakin domain mutations arrested elongation for similar reasons, we examined the distribution of LET-805::GFP and MUP-4::GFP, the basal and apical hemidesmosome receptors, respectively. For this, we generated knock-in versions of MUP-4::GFP and LET-805::GFP. After PAK-1 knockdown in the *vab-10A*(Δ*SH3*)::*mCherry* or *vab-10*(Δ*SR8*) backgrounds (Fig. 4A), basal LET-805::GFP hemidesmosomes were normal until the 2-fold stage, and then displayed integrity defects mainly in the region where embryo curvature is the highest (Fig. 4A and Movies S4-S5; see arrows and dotted line in Fig. 4A). Specifically, the LET-805::GFP signal appeared to detach from the outer body wall and to collapse internally as if not held anymore in *vab-10*(Δ*SH3*), consistent with the observation that muscles also detached from the body wall (Fig. S2). The basal VAB-10(mc62)::mCherry signal remained associated with it, but was much dimmer, suggesting that detachment occurred within the epidermis layer (Fig. 4B, bottom three rows and Movie S6). By contrast, at the apical side, the signal appeared intense although often less compact (Fig. 4B). These defects, which were not observed in VAB-10A(+)::mCherry control embryos after PAK-1 knockdown (Fig. 4B upper row), are strongly reminiscent of those previously observed in *vab-10A*(*e698*); *pak-1*(*ok448*) double mutants (Zhang et al., 2011). Interestingly, the same LET-805::GFP detachment phenotype was observed in *vab-10*(Δ*SR5h1-h2*) embryos (Fig. 4C); however, in contrast to the situation observed in embryos homozygous for the presumptive null allele *vab-10*(*h1356*) (Bosher et al., 2003), LET-805 could still be observed in the remainder of the embryo arguing that *vab-10*(Δ*SR5h1-h2*) is only a hypomorph. The apical hemidesmosome receptor MUP-4::GFP was also affected at the turn of the embryos and displayed a much fainter intensity (Fig. 4D). These results and the aforementioned genetic interactions raise the hypothesis that the SH3 and SR8 domains within the VAB-10A/plectin isoform together with PAK-1 are acting in the mechanotransduction process to allow hemidesmosomes to sustain tension beyond the 2-fold stage. This does not preclude an additional role of these domains within the VAB-10B/MACF isoform. We suggest that hemidesmosomes preferentially rupture in the most convex part of the embryo because tension is higher where the curvature is higher.

**Figure 4.**
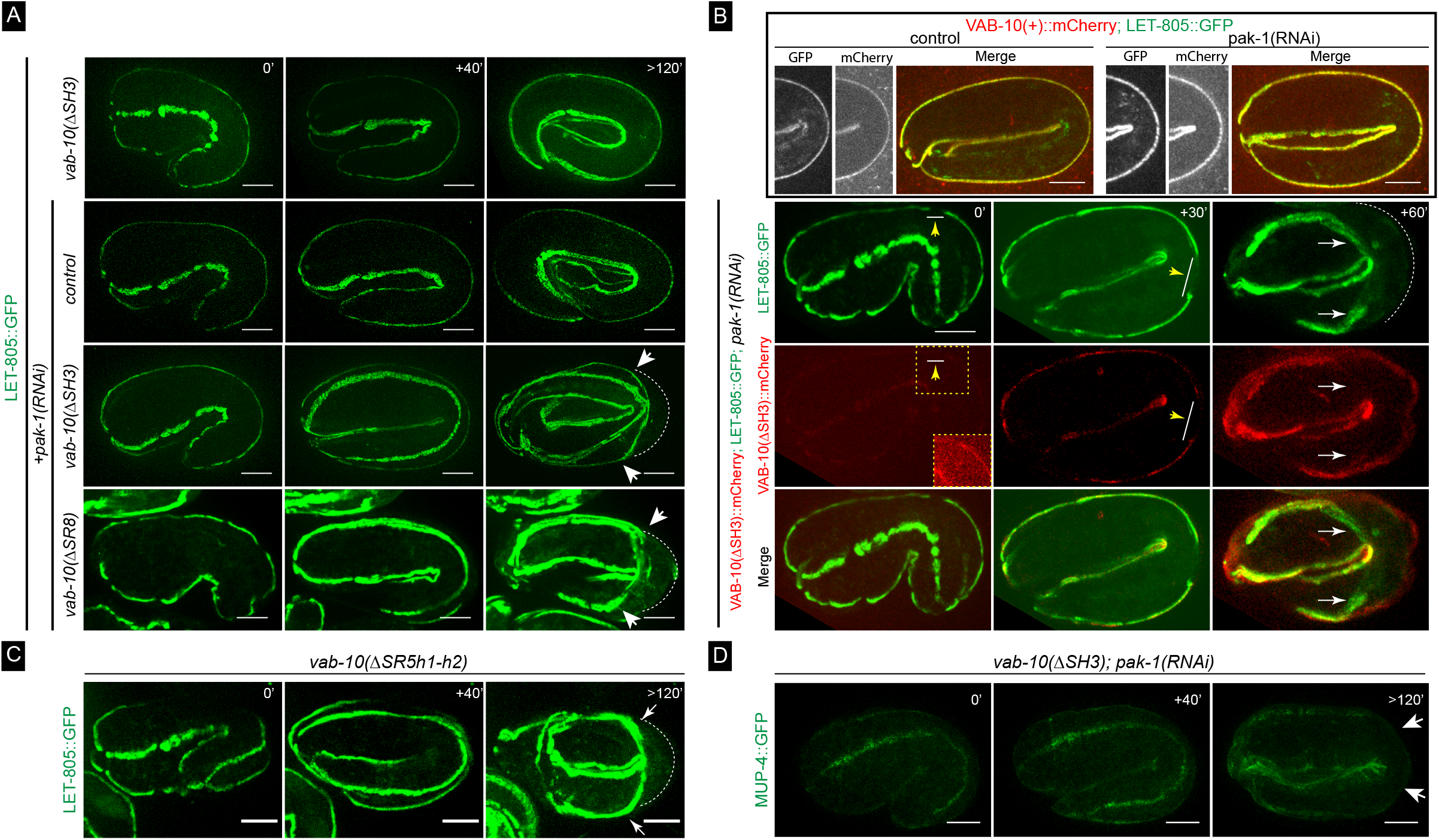
The maintenance of hemidesmosome integrity requires multiple plakin sub-domains. Spinning disc fluorescence micrographs from time-lapse movies of embryos; each panel represents the projection of approximately 10-15 focal planes (out of generally 60). **(A)** Embryos homozygous for the CRISPR knockin LET-805::GFP (*mc73*; N=35) with the genotype indicated on the left. Note that in the *vab-10*(Δ*SH3*); *pak-1*(*RNAi*) embryo (3^rd^ row) and the *vab-10*(Δ*SR8*); *pak-1*(*RNAi*) embryo (4^th^ row), LET-805::GFP signal has detached from the body wall on the convex side and collapsed internally (white arrows, area of detachment; white dotted line, body wall - compare to 1^st^ and 2^nd^ rows). Pictures in rows 2-3 are from Movies S4 and S5, respectively. **(B - row 1)** Wild-type VAB-10(+)::mCherry; LET-805::GFP embryo alone or after *pak-1*(*RNAi*); in both cases the 1^st^ panel shows the GFP signal in the turn of the embryo, the 2^nd^ the mCherry channel and the 3^rd^ the merge channel at the 2.1/2.2-fold stage. **(B - rows 2-4)** Homozygous for *vab-10*(*mc62*)::*mCherry*; *LET-805*::*GFP*(*mc73*), *pak-1*(*RNAi*) embryo at three different stages (last being 2.2-fold stage equivalent). This embryo displayed a small interruption in the hemidesmosome line visible at earlier stages (arrow and line), which prefigured the area of detachment (white arrows); 20 out of 21 embryos showed detachment in the convex part, among which 7 had a detectable interruption. 2^nd^ row, LET-805::GFP signal; 3^rd^ row, VAB-10(mc62)::mCherry signal (the inset corresponds to a small area for which the intensity was increased with FiJi); 4^th^ row, merge. Pictures in row 4 are from Movie S6. **(C)** Embryo homozygous for *vab-10*(Δ*SR5h1-h2*); *LET-805*::*GFP*(*mc73*) (all embryos showed this phenotype; N=15). Note a detachment as in panel A rows 3-4 (arrows). **(D)** *vab-10*(Δ*SH3*); *pak-1*(*RNAi*) embryo homozygous for MUP-4::GFP knockin (*mc121*; N=30). The MUP-4::GFP signal is much weaker on the convex side (between white arrows). All examined embryos showed a detachment. Scale bars, 10 μm.

### GIT-1 recruitment to hemidesmosomes depends on VAB-10 and LET-805

We previously established that GIT-1 recruitment to hemidesmosomes is the first detectable step in the mechanotransduction pathway (Fig. 1C). A possibility could be that a hemidesmosome-associated protein (or proteins) acting as a mechanosensor able to sense changes in tension when muscles contract and relax that recruits GIT-1 (Fig. 1C). To test which hemidesmosome component would be involved in recruiting GIT-1, and more specifically to explore if the plakin domain could do so, we examined whether specific plakin sub-domains deletions or hemidesmosome component depletion affect GIT-1 distribution.

In contrast to the severe reduction of GIT-1::GFP signal observed in embryos defective for the essential muscle protein UNC-112 (Zhang et al., 2011) (Fig. 5A, bottom row), we found that absence of the SH3 domain did not affect the GIT-1::GFP signal (Fig. 5A, 2^nd^ row). By contrast, absence of the first two helices of the SR5 domain partially reduced GIT-1 recruitment and strongly compromised hemidesmosome integrity where embryo curvature is the highest (Fig. 6A, line *vab-10*(Δ*SR5h1-h2*)), consistent with the global hemidesmosome detachment at that position (see Fig. 4C). Since some GIT-1 remained at hemidesmosomes in *vab-10*(Δ*SR5h1-h2*) mutants, we tested if other hemidesmosome proteins help recruit GIT-1 to hemidesmosomes by comparing the intensity and continuity along the anterior-posterior axis of the GIT-1 signal (Fig 5B-B’) after RNAi-induced depletion of LET-805, VAB-10 or PAT-12, three essential hemidesmosomal components (Zhang and Labouesse, 2010). We found that both parameters were reduced with the following grading in severity: *unc-112*(*RNAi*) > *let-805*(*RNAi*) ≥ *vab-10*(*RNAi*) ≈ *vab-10*(Δ*SR5h1-h2*) ≥ *pat-12*(*RNAi*) (Fig. 5C-D). In particular, a strong VAB-10 RNAi knockdown and the deletion of the first two helices of the SR5 domain resulted in very similar phenotypes, whereas *pat-12*(*RNAi*) moderately affected GIT-1::GFP levels and continuity, while still inducing a strong hemidesmosomal detachment defect (Fig. 5, see time +60 min). Importantly, loss of LET-805 does not significantly reduce VAB-10A levels (Hresko et al., 1999), suggesting that LET-805 might be directly involved in recruiting GIT-1. We conclude that GIT-1 recruitment is likely to involve LET-805 and potentially the VAB-10 SR5 domain.

To examine through another approach whether GIT-1 is indeed in close proximity with VAB-10, we used a bi-molecular fluorescence complementation (BiFC) strategy (Hu et al., 2002) (Fig. S3A). We generated CRISPR knock-in strains expressing GIT-1 linked at the C-terminus to the 173 first residues of Venus, and VAB-10A linked at the C-terminus to the last 83 residues of Venus. As a positive control, we co-expressed the same GIT-1::Venus(1-173) and a CRISPR-generated PIX-1::Venus(155-238), since vertebrate Git1 and β-PIX form a complex (Frank and Hansen, 2008). We found that co-expressing GIT-1::Venus(1-173) with PIX-1::Venus(155-238) or VAB-10A::Venus(155-238) gave a clear hemidesmosomal signal (Fig. S3B), which was similar to that of endogenous VAB-10A and VAB-10B, and of a novel CRISPR knock-in of GIT-1::GFP (Fig. S3C-D). By contrast, embryos expressing only one Venus moiety failed to give a signal (Fig. S3B). We conclude that VAB-10A and GIT-1 are located within less than 5-10 nm, the maximum distance beyond which a BiFC signal cannot be detected (Ciruela et al., 2010; Hu et al., 2002). Since the fluorophore moieties were at the C-terminus of each protein, and since VAB-10A is a large protein of 3400 residues, additional methods will be required to define if GIT-1 directly interacts with VAB-10.

**Figure 5.**
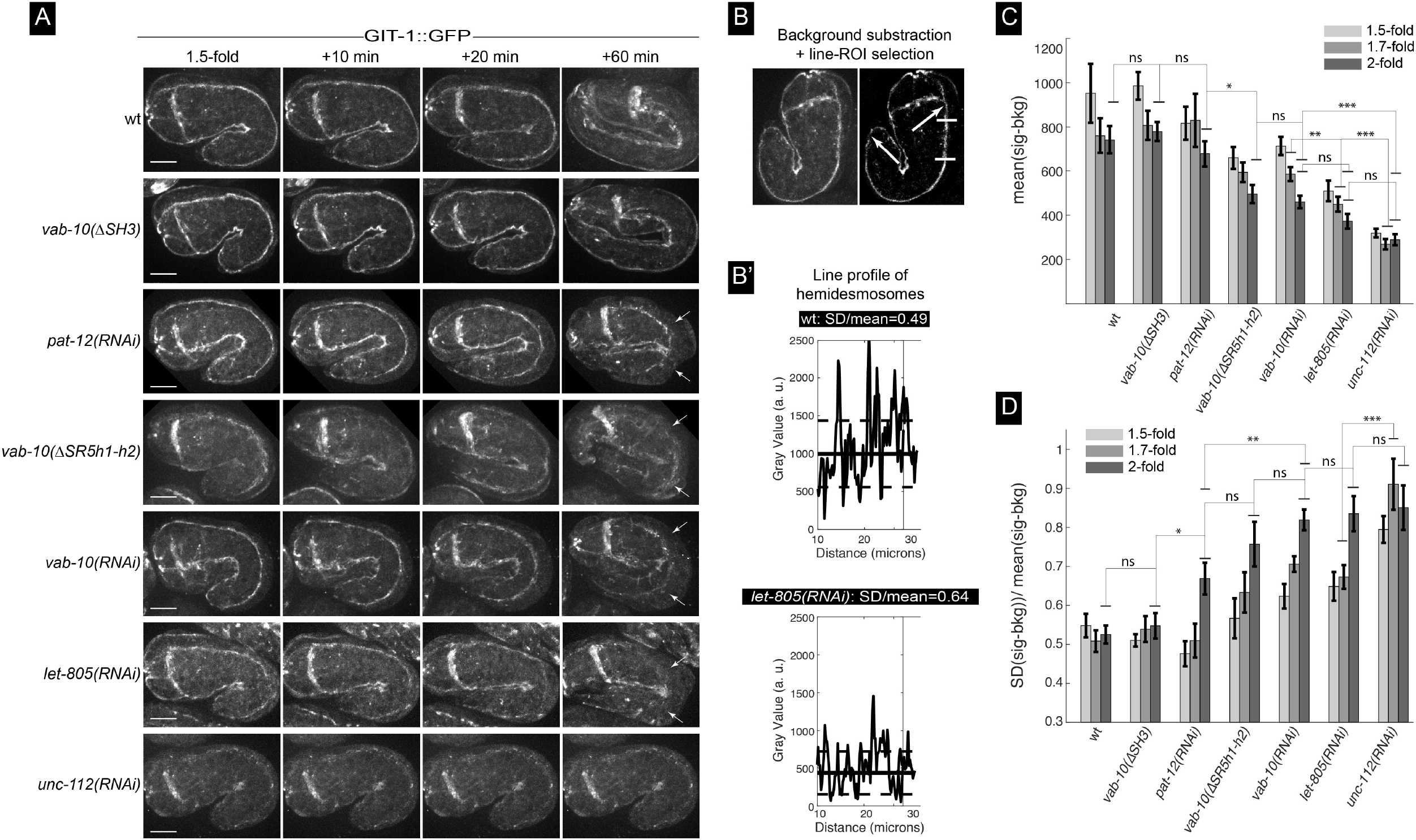
Differential requirement of hemidesmosome components for GIT-1 recruitment. **(A)** Spinning disc fluorescence pictures from movies S7 showing the distribution of the GIT-1::GFP knockin (allele *mc86*) at the 1.5-fold stage and 10, 20 and 60 minutes later for the genetic backgrounds indicated on the left. Each picture shows only the top two hemidesmosomes. Note how the hemidesmosome signal is collapsing internally from the most convex side of the embryo for *pat-12*, *vab-10* and *let-805* knockout embryos as well as for *vab-10*(Δ*SR5h1-h2*) embryos (arrows). **(B-B’)** Procedure used for image analysis to quantify the signal intensity and continuity: after background subtraction (**B**), the signal intensity was measured along the hemidesmosome signal between the two arrows. (**B’**) Examples of the signal for two backgrounds, but just for the area bracketed by short horizontal white lines in (**B**). **(C-D)** Signal intensity (**C**) and continuity (**D**) for the backgrounds illustrated in (**A**) at three time points. Scale bar, 10 μm. Statistical tests: ns, not significant; *, p < 0.5; **, p< 0.01; ***, p< 0.001.

### Biophysical evidence that the SH3 domain can be mechano-sensitive

The previous sections establish that the region shielding the SH3 domain of VAB-10 is essential for mechanotransduction, but that the SH3 domain is unlikely to directly interact with GIT-1. To further explore how this region responds to force, we used Molecular Dynamics (MD) simulations. Our previous MD simulations of the homologous plakin domains of desmoplakin and plectin under a stretching force have suggested that the SH3 domain mechanically stabilizes the spectrin repeats, and that force relieves the auto-inhibition of the SH3 domain by the preceding SR4 domain (Daday et al., 2017). It predicted that the SH3 domain could have a mechanosensing role.

We performed the same simulations for a homology model of VAB-10 as shown in Fig. 1E (see Methods). Throughout our equilibrium MD simulation, the contact area between the SH3 domain and the rest of the protein was stable, with quartiles 10.3-11.5 nm^2^. This indicates that the VAB-10 SH3 domain interacts with the spectrin repeats in a very similar manner as the previously studied homologue (Daday et al., 2017). To examine the possibility that the SH3 domain might be mechano-sensitive, we subjected 10 snapshots from our equilibrium simulations to stretching forces acting on the termini in subsequent force-probe MD simulations. Similar to the tendencies observed for plectin, we find that the VAB-10 spectrin repeats always unfolded before the SH3 domain, i.e. the SH3 domain was invariably exposed and activated before its unfolding (Fig. S4A-B). This activation could happen at the very first stage of the unfolding of the spectrin region, or in the second stage; hence VAB-10 behaves like plectin and desmoplakin, which we examined in previous simulations (Fig. S4C). In the case of plectin, early activation, that is, after partial unfolding of at most one spectrin repeat, happened in about ⅕ of the simulations, while in the case of VAB-10, this happened even more frequently in ⅓ cases (Fig. 6A). We also computationally analyzed the plakin domain of VAB-10 carrying the Δ*SR5h1-h2* deletion (mc64), which removes the first two helices of the SR5 spectrin repeat shielding the SH3 domain. Our model predicts the SH3 domain to still bind to the upstream SR4 (Fig. S4D), suggesting that the phenotype induced by this deletion is unlikely to result from the permanent availability of the SH3 domain to bind other proteins, and thereby act as a constitutively active protein. Taken together, our results are compatible with the notion the VAB-10 SH3 domain can stabilize its plakin domain and potentially bind to other proteins if tension is exerted on the plakin region.

**Figure 6.**
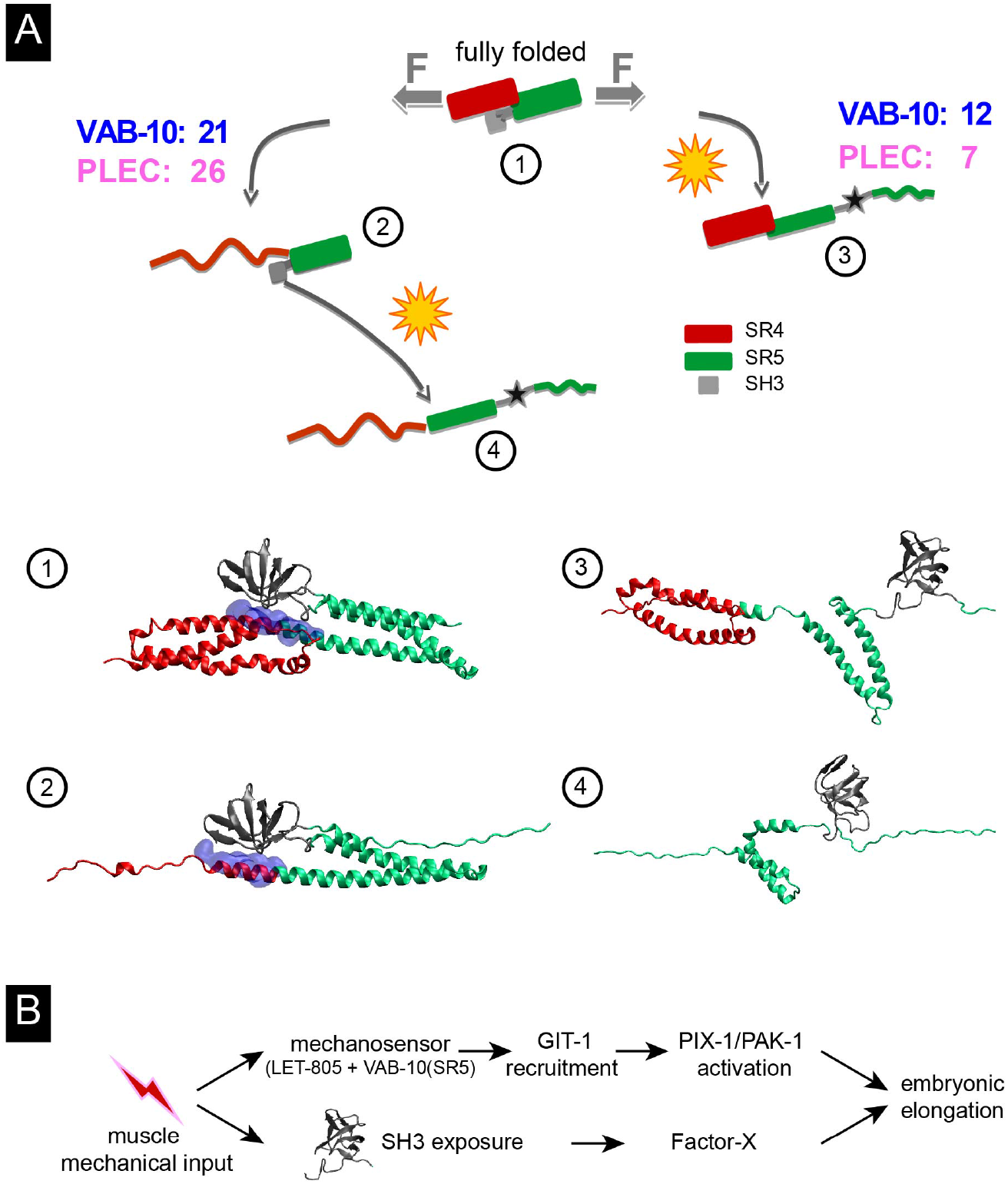
Unfolding forces exerted on the plakin domain of VAB-10 unmask the SH3 domain. **(A)** Mechanical unfolding pathways of VAB-10 and plectin lead to SH3 activation. The numbers indicate how many trajectories showed SR4 unfolding first (left) versus SR5 unfolding first (right). Numbers for plectin are based on (Daday et al., 2017). Red, SR4; green, SR5 (the thickness of the green rectangle is proportional to the number of helices contributing to it); blue, SH3-SR4 interface; grey: SH3, star: exposure; wavy red/green lines, unfolded SR helices. **(B)** Model for mechanotransduction at the hemidesmosome (see text).

## Discussion

This study combines molecular genetic analysis of the *C. elegans* spectraplakin VAB-10/plectin with Molecular Dynamics simulation to functionally test the function of its plakin domain. Our data establish that the SH3 domain with its shielding SR5 spectrin repeat, and the SR8 domain are essential to enable mechanotransduction at hemidesmosomes.

The recent crystal structure of the plakin domain of plectin revealed that this domain composed of nine spectrin repeats should adopt an extended rod-like shape (Ortega et al., 2011; Ortega et al., 2016). Intriguingly, it also revealed the presence of an SH3 domain embedded within the central spectrin repeat (SR5). This SH3 is atypical inasmuch it is missing several residues that normally ensure the interaction with Pro-rich target sequences within prototypical SH3 domains. Instead, it exhibits multiple interactions with the previous spectrin repeat SR4, questioning the notion it could act as a *bona fide* SH3 domain, without ruling out that it could interact with proteins in a non-canonical way (Ortega et al., 2011). By using the power of *C. elegans* molecular genetic tools and CRISPR-mediated recombination approaches, we could test the function of the VAB-10 plakin domain. Our key findings are that removal of the first two helices of the SR5 domain upstream of the SH3 domain induced highly penetrant embryonic elongation defects, but that deletion or point mutations within the SH3 domain as well as deletion of the SR7 or SR8 domain barely affected VAB-10 function on their own. Furthermore, alterations of the SH3 domain or a SR8 deletion, but less frequently of a SR7 deletion, induced embryonic elongation defects when combined with loss of the signaling proteins GIT-1 or PAK-1. The specificity of their phenotypes, and the observation that *vab-10*(Δ*SR5h1-h2*) deletion did not affect as much LET-805 distribution as the presumptive null allele *vab-10*(*h1356*), strongly argues that these deletions do not induce major structural defects in VAB-10 polypeptides but modify distinctive aspects of VAB-10 function.

The proteins GIT-1, PIX-1 and PAK-1 form the core of a mechanotransduction process induced by muscles in the epidermis to maintain hemidesmosome integrity when tension rises during elongation (Zhang et al., 2011). Genetically speaking, the synergistic effects observed between *vab-10*(Δ*SH3*) or *vab-10*(Δ*SR8*) deletions and *pak-1* or *git-1* mutations implies that the SH3 and SR8 domains are acting in a parallel rather than a linear pathway with GIT-1/PAK-1 in the process of hemidesmosome maintenance. Furthermore, these data predict that an unidentified Factor-X acts in parallel to GIT-1/PIX-1/PAK-1 downstream of the mechanical input and the SH3 domain, which might either recruit or activate Factor-X (Fig. 6B). Finally, these genetic data also imply that neither the SH3 nor the SR8 domain is involved in recruiting GIT-1 to hemidesmosomes, consistent with our observations that GIT-1 was still present at hemidesmosomes in the absence of the SH3 domain. Instead, two arguments suggest that GIT-1 is recruited to hemidesmosomes at least in part through the interface between the SR5 domain and the presumptive hemidesmosome receptor LET-805. First, *vab-10*(Δ*SR5h1-h2*) deletion and to a higher degree *let-805*(*RNAi*) reduced GIT-1 level at hemidesmosomes. Second, their effect are unlikely to be indirect since LET-805::GFP levels remained high in *vab-10*(Δ*SR5h1-h2*) embryos (Fig. 4C), and since a *let-805* mutation does not reduce VAB-10A levels as monitored with the MH5 antibody against VAB-10A (Hresko et al., 1999). Our unpublished yeast 2-hybrid screens taking various parts of GIT-1 as baits, or conversely taking either the SR5 or the SH3 domains as baits, failed to identify any significant prey; although such negative results should be taken with care, our failure might stem from the fact that the SR5-SH3 region as well as GIT-1 interact only as a multi-protein complex that cannot be picked up through yeast 2-hybrid screens. A key objective for future studies will be to identify the predicted Factor-X and the contact points between GIT-1 and hemidesmosome components.

How would proteins contributing to the mechanotransduction process assemble, and would they form a stable or a dynamic complex? Our Molecular Dynamics simulations predicted that tension should expose the SH3 domain due to the unfolding of the SR4 and/or SR5 domains, as previously predicted for plectin and desmoplakin (Daday et al., 2017). Since muscles repeatedly contract and relax (Lardennois et al., 2019; Zhang et al., 2011), one can speculate about various and not necessarily exclusive scenarios. One possibility could be that a multi-protein signaling complex including GIT-1 and Factor-X stably assembles along the VAB-10 plakin region once muscles contract; such a complex would fail to form in the absence of the SR5, SH3 or SR8 domains. Another possibility could be that the cyclic muscle-induced tension pattern promotes the transient and periodic recruitment of GIT-1 to the SR4-SR5 domains either in their folded or unfolded configuration. A third possibility would be that the SH3 domain recruits Factor-X depending on tension, or in a more complex version that different signaling complexes alternatively bind and unbind to the SR4-bound SH3 and to the free SH3 to facilitate hemidesmosome remodeling. Interestingly, the SH3 domain of the plectin-1c isoform alone or more strongly in a presumptive SR4-SR5 folded configuration can bind microtubule-associated proteins to destabilize their interaction with microtubules (Valencia et al., 2013), implying that the SR4-SR5-SH3 region and the SH3 alone can interact with other proteins. Interestingly, we previously reported that microtubule depletion in the *vab-10A*(*e698*) background leads to a phenotype very similar to that observed in the mutants described herein (Quintin et al., 2016).

The tension-dependent recruitment of GIT-1 and/or Factor-X in some of these scenarios would posit VAB-10 as a mechanosensor. The paradigm for mechanosensing is largely based on talin and α-catenin, the two best-characterized mechanosensors. In both cases, it requires the unfolding of talin and α-catenin internal domains following their interaction with a junction receptor and with actin through terminal domains (del Rio et al., 2009; Hu et al., 2017; le Duc et al., 2010; Yao et al., 2016; Yao et al., 2014; Yonemura et al., 2010). There are two intriguing similarities between talin and plakins. One is that the plakin domain, much like the central part of talin, consists in several repeated domains (9 spectrin repeats versus 13 alpha helix bundles, respectively) that can be unfolded upon tension (Law et al., 2003; Lenne et al., 2000; Yao et al., 2016). Another is that among the bundles composing talin, the central repeat (R8) loops out of the preceding R7 and unfolds with it, which is partially reminiscent of the SH3 domain being inserted within the central SR5 domain of the plakin domain (Yao et al., 2016). Moreover, the Rho GTPase Activating Protein DLC1 (Deleted in Colorectal Cancer1) is active when bound to the folded R8 but inactive and unbound when R8 unfolds (Haining et al., 2018). These features may characterize ECM-linked mechanosensors.

As recalled above, mechanosensing through talin requires it to bind to integrin and actin. In the case of VAB-10A, the situation would be conceptually similar although molecularly presumably different. As illustrated in Fig. 1B, there are two hemidesmosome-like structures in the epidermis, one basally and one apically, each associated with a different ECM. Both hemidesmosomes with their bridging intermediate filaments act as tendons linking muscles to the apical ECM. In the 3D space of embryos, when muscles contract in the anterior-posterior direction, this tendon-like structure oriented radially will come under high tension. Although not yet biochemically confirmed, genetic experiments suggest that VAB-10A should bind to the basal transmembrane receptor LET-805 (Bosher et al., 2003; Hresko et al., 1999; Zhang and Labouesse, 2010), much like plectin binds to β4-integrin through its two calponin-homology domains and probably the SR4-SR5-SH3 region (Frijns et al., 2012; Koster et al., 2004). It is likewise likely that VAB-10A can bind to MUP-4 on the apical side. While LET-805 basally and MUP-4 apically bear no homology to β4-integrin, their cytoplasmic tails are long enough to create multiple binding surfaces with VAB-10. Hence, when muscles exert tension on hemidesmosomes, the spectrin repeats of the VAB-10 plakin domain could become unfolded to expose its SH3 domain.

In conclusion, our work reveals the key role of the VAB-10 plakin domain in mediating mechanotransduction in vivo, and possibly mechanosensing. Molecular Dynamics simulations suggest that the SH3 domain can alternate between an SR4-interacting state and a free state, with the latter being induced by force. Genetic analysis of *C. elegans* embryos suggests that the situation in vivo is complex and could involve multiple protein complexes acting in parallel. By extension, our data suggest that the SH3 domain and its shielding region are likely to play a critical signaling role in vivo in other spectraplakins too.

## Materials and Methods

### Strains and culture

*Caenorhabditis elegans* wild-type strain and transgenic animals derived from the *N2* strain. They were used for all experiments and maintained as described in (Brenner, 1974). A complete list of strains and associated genotypes used in this study are included in supplementary Table 1.

### Molecular Biology

Plasmid constructions were performed using Standard PCR method (Barstead et al. 1991) and general molecular biological techniques used as described in Sambrook et al. (1989). To construct knock-in DNA plasmid-based repair templates used for CRISPR/Cas9-mediated genome editing, we amplified the >1.5Kb upstream and >500bp downstream sequences from the *N2* genomic DNA, and the fluorophore-encoding fragment from pre-existing vectors by using Phusion DNA polymerase. PCR fragments were analyzed on agarose gel and concentrated by either PCR clean-up or gel purification kit (QIAGEN) and their concentrations measured using a Nanodrop (Eppendorf) before final assembly with pJET1.2 vector using NEBuilder HiFi DNA Assembly Cloning Kit (New England Biolabs). Small Guide encoding plasmids for CRISPR were generated by an overlap extension PCR method performed on template pML2840.

To avoid Cas-9 cleavage of the homologous repair template, the Cas9 site of the repair template was modified by introducing synonymous mutations either directly into the primers used for fragment amplification or separately by Q5 site directed mutagenesis kit (New England Biolabs). All DNA plasmids used for genome editing were transformed into DH5α competent cells and subsequently purified by miniprep (PureLink Quick Plasmid Miniprep Kit (Thermo Fisher Scientific) or Plasmid Midi Kit (QIAGEN). All final DNA constructs were sequence verified before use. The sequences of sg-primers (SIGMA) used in this study are included in Supplementary Table 2.

### CRISPR/Cas9-mediated genome editing

Wild type *N2 C. elegans* genetic background was used to generate *vab-10* CRISPR/Cas9 alleles; *let-805* and *mup-4* GFP knock-in strains were generated after injection in an *unc-119*(*ed3*) background (Dickinson et al., 2013). The sgRNA plasmid, knock-in repair template plasmid, Cas9 encoding plasmid and the appropriate co-injection marker (PRF4/ myo-2::mCherry/ myo-2::GFP) were co-injected into *N2* animals. Typically, injection mixes were prepared in DNase and RNase free MilliQ water and contained a combination of 50-100 ng/μl sgRNA plasmid (targeting specific gene), either 50 ng/μl repair template plasmid or 20-50 ng/μl ssODNA (PAGE-purified oligonucleotide) repair template, and co-injection markers pRF4 [rol-6 (su1006)]100 ng/μl, 2.5 ng/μl myo-2p::mCherry/ myo-2p::GFP. Injection mixes were spun down in a microcentrifuge (Eppendorf) for at least 10-30 minutes at 14,000 RPM prior to use. 30-40 young adult hermaphrodites were injected in the germline using an inverted micro-injection set up (Eppendorf). After injection, one animal per NGM food plate was dispatched and grown at 20 °C for 2-3 days. F1 animals carrying co-injection markers were picked and singled out on separate NGM OP50 plate, and grown at 20 °C until eggs or larvae were spotted. Each F1 mother was lysed in separate PCR-tube and standard worm PCR protocol was followed using appropriate pairs of primers (annealing in the inserted sequence and a genomic region not included in the repair template). For the construction of some transgenic strains, we also used Co-CRISPR the *dpy-10* phenotype (Paix et al.,2015) or integration of a self-excisable cassette carrying a visible marker (Dickinson et al., 2015) method. All genotyping experiments were carried out using standard worm PCR method. Confirmed alleles were sequenced and verified (Eurofins).

### RNA-mediated interference (RNAi)

RNAi experiments were performed either by feeding on *HT115 Escherichia coli* bacteria strains generating double-stranded RNA (dsRNA) targeting genes of interest or by injection of in-vitro synthesized double-stranded RNA on young L4 stage of the animals. Feeding RNAi clones for *crt-1* and *mec-8* were used from the Ahringer-MRC feeding RNA interference (RNAi) library (Kamath et al., 2003). RNAi feeding was performed using standard procedures, with 100 μg ml^−1^ ampicillin/1 mM IPTG (Sigma). Empty L4440 RNAi vector served as a control. Other experiments involving RNAi were done by injection of dsRNA.

To generate dsRNA for the injection, genomic fragments were PCR amplified using Phusion DNA polymerase (ThermoFisher) and these fragments were further served as templates for in-vitro dsRNA synthesis using T3 or T7 mMESSAGE mMACHINE Kit (Ambion, USA). For gene knockdown experiments by feeding, L4/L1 hermaphrodites were grown on RNAi plates for 24-36 hrs; for dsRNA injection, 20-30 young L4 hermaphrodites were injected with dsRNA targeting gene of interests and grown for 14-20 hrs prior to experiments.

### Spinning disk and Nomarski microscopy

For live imaging, embryos were picked from NGM plate by mouth pipette, washed thoroughly in M9 medium and mounted on 2-5% agarose pads after sealing the slides with paraffin oil. Spinning disk imaging of embryos was performed using a Roper Scientific spinning disk system (Zeiss Axio Observer Z1 microscope, Yokogawa CSUX1-A1 spinning disk confocal head, Photometrics Evolve 512 EMCCD camera, Metamorph software) equipped with a 63X, and 100X oil-immersion objective, NA=1.4. The temperature of the microscopy room was maintained at 20 °C. Images of embryos were acquired in either streaming mode with 100 ms exposure, or timelapse mode with 100 ms exposure and 5 minutes intervals. Laser power and exposure times were kept constant throughout the experiments for specific strains and their control genotypes. For the quantification of hemidesmosomes, images were acquired with 100 ms exposure time in stream mode with 0.3 μm step size. Images were processed and quantified using FIJI. Fluorophores used in this study include eGFP, GFP (65C), mCherry and split VENUS.

Time-lapse DIC movies were acquired using a Leica DMI6000 upright microscope, a 40X or 63x oil-immersion objective NA=1.25 and a Photometrics Coolsnap HQ2 camera (Photometrics, AZ, US) placed in the temperature-controlled room at 20 °C. For *C. elegans* larvae image acquisition, animals were anesthetized using 0.1 mmol/L levamisole in M9 buffer and mounted on 2% agarose pad.

### Homology modelling

We used MODELLER (Sali and Blundell, 1993), version 9.14, through the interface in UCSF Chimera version 1.12 (Pettersen et al., 2004), for homology modelling. Currently, two crystal structures of proteins with SH3 insertions are available: 3PE0 (plectin) and 3R6N (desmoplakin). The main difference between these two structures is the presence of a small helix, B0, in spectrin repeat 5. Given that a later investigation found no such small helix in plectin (Ortega et al., 2016), we chose to model VAB-10 using 3R6N (desmoplakin), despite the fact that the sequence identity is lower (27.37% for desmoplakin as opposed to 32.10%). When compared to 3R6N, the obtained model has a distance of 0.46 **Å** between the 265 pairs of Cα atoms within 2.0 **Å** of each other out of a total of 274 pairs. As given by MODELLER’s estimates, the homology model is at an RMSD of 2.21 **Å** and an overlap of Cα atoms of 0.91 within 3.5 **Å**. We compared the three structures (3PE0, 3R6N, and our obtained homology model) in Fig. 1E. We also obtained homology models for the mutant protein VAB-10(mc64) lacking the first two helices of SR5, which show the SH3 domain still associated to helix 2C, but helix 3C is shown to loop back onto the rest of the structure in two different conformations (Fig. S3D).

### Molecular dynamics simulations

We followed the same protocol as described in our previous work (Daday et al., 2017). In short, we used the Amber99SB-ILDN* force field (Liu et al., 2016) with a TIP3P water model (Jorgensen et al., 1983) and virtual sites(Berendsen and van Gunsteren, 1984), allowing for a 5 fs time step. All MD simulations were performed using GROMACS, version 5.0 (Pronk et al., 2013). We performed equilibrium molecular dynamics on approximately 190k atoms in a dodecahedron box for 1 μs, and thereafter we chose the top 10 frames through cluster analysis between 100-1000 ns, with a cutoff of 0.088 nm. These 10 frames were later used for force-probe simulations. 10 simulations were performed at each of the velocities 1, 1/3, 1/10 nm/ns, and 3 simulations were performed at the velocity of 1/30 nm/ns. All other parameters were identical to the procedure described in our previous work, in particular for the re-solvation procedure for plectin. The equilibrium molecular dynamics simulation shows a backbone RMSD with an inter-quantile range between 3.4-4.4 **Å**. The helicity of the construct is very stable, with quartiles at 209 and 214 residues out of the 274 in our structure, and the β-strand content is stable, with quartiles at 35 and 40 residues and a small positive drift (1 more residue in strands every 236 ns). Overall, we consider the homology model to be a good representation of this region of VAB-10.

### Image processing, quantification and statistical analysis

Images in Fig. 5 were analyzed as follows: the background was estimated by Gaussian filter of the original image with a width of 30 pixel; the background image was subtracted from the original one; the line profile of the GIT-1 signal along the hemidesmosomes was measured on the subtracted image. We recorded two observables from the line profile: the average signal and its standard deviation. The average signal is shown in Fig. 5C, whereas the ratio between standard deviation and average signal (coefficient of variation) is shown in Fig. 5D as a measure of the discontinuity. Indeed a fragmented signal shows higher relative fluctuations. All images were analysed using the ImageJ (FiJi) software (NIH, Bethesda, Maryland, USA; http://rsb.info.nih.gov/ij/) and statistical test were performed using MATLAB R2018b (The MathWorks Inc., Natick, MA). For Fig. 5 a Wilcoxon test was used, whereas for Fig. 2J a chi-square test was applied.

## Supporting information

folder with seven supplementary movies

## Acknowledgements and Funding

We thank the Caenorhabditis Genetics Center (funded by the NIH Office of Research Infrastructure Programs P40 OD010440) for strains, and the IBPS Imaging Facility for advice. This work was supported by a European Research Council (grant #294744), ANR (ANR-11-BSV2-0023) and installation funds from the Centre National de la Recherche Scientifique (CNRS) and University Pierre et Marie Curie (UPMC) to ML. This work was also made possible by institutional funds from the CNRS, University of Strasbourg and UPMC, the grant ANR-10-LABX-0030-INRT which is a French State fund managed by the Agence Nationale de la Recherche under the framework programme Investissements d’Avenir labelled ANR-10-IDEX-0002-02 to the IGBMC. FG acknowledges funding from the Klaus Tschira Foundation, from the German Research Foundation (DFG) through the priority programme SPP1782, and from the state of Baden-Württemberg and the DFG through grant INST 35/1134-1 FUGG.

---

**Figure S1.**
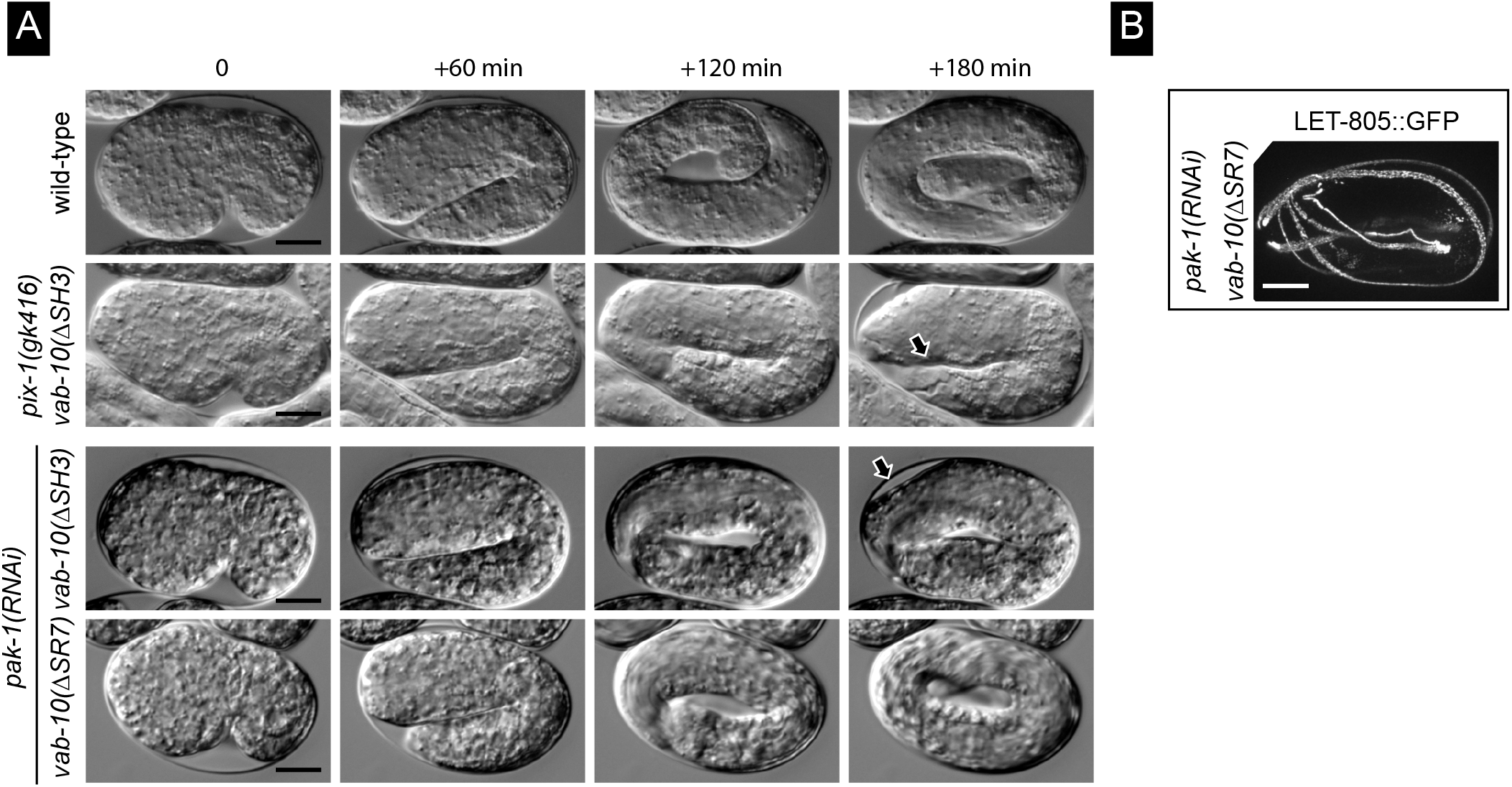
Genetic interactions between the *pak-1* and *vab-10*(Δ*SR7*) or *vab-10*(Δ*SR8*) deletions. **(A)** DIC micrographs taken from time-lapse movies of wild-type (1^st^ row), *vab-10*(Δ*SH3*); *pix-1*(*gk416*) (2^nd^ row), *vab-10*(Δ*SH3*); *pak-1*(*RNAi*) (3^rd^ row), *vab-10*(Δ*SR7*); *pak-1*(*RNAi*) (4^th^ row). **(B)** Homozygous *vab-10*(Δ*SR7*) (1^st^ row) and *vab-10*(Δ*SR8*) (2^nd^ row) carrying the LET-805::GFP (*mc73*) after RNAi against *pak-1*. Note that hemidesmosomes collapsed in the turn of the *vab-10*(Δ*SR8*) embryo. Scale bars, 10 μm.

**Figure S2.**
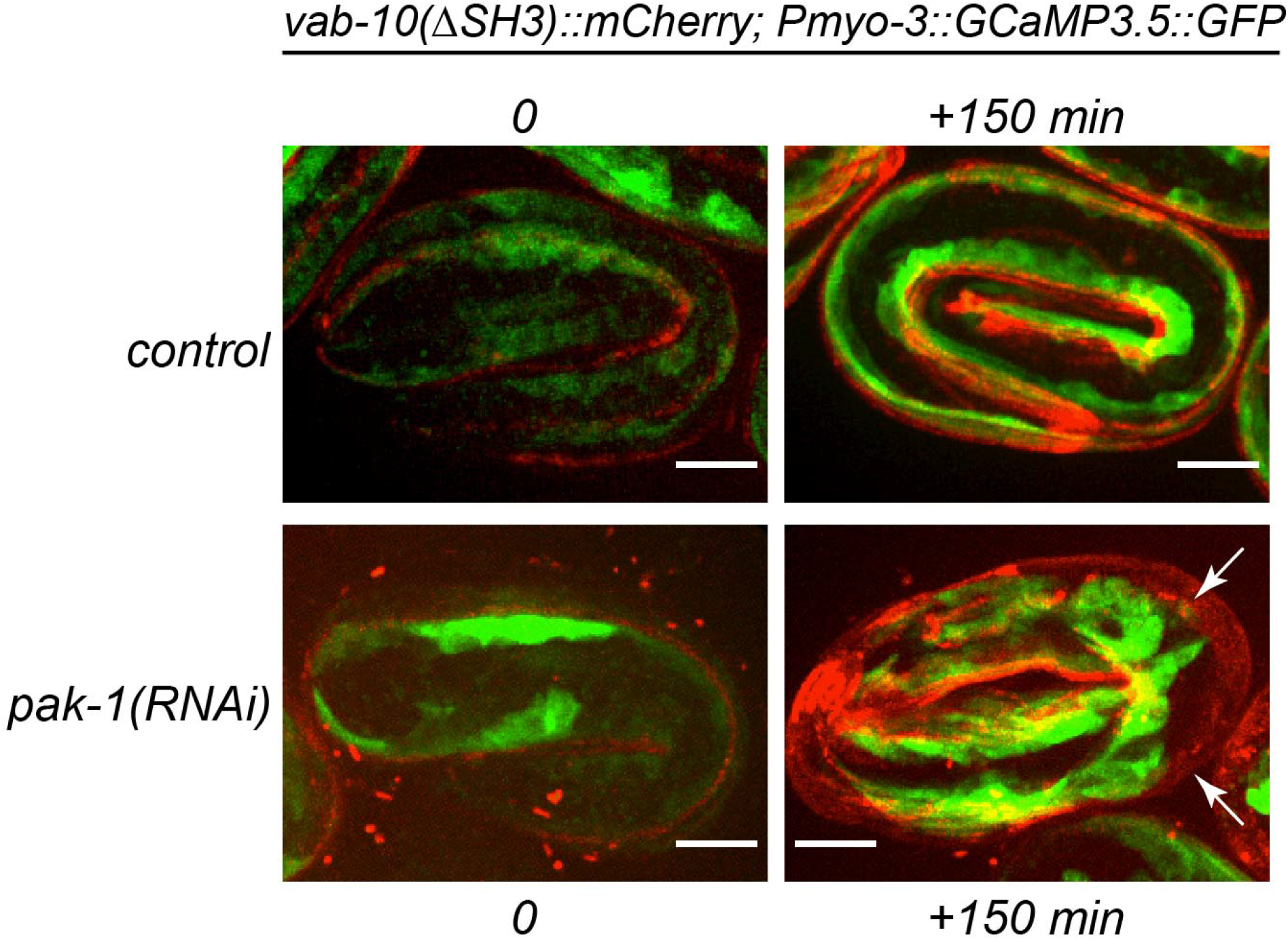
Hemidesmosome defects trigger muscle detachment. Fluorescence pictures taken from a video from homozygous *vab-10*(Δ*SH3*)::*mCherry* carrying a transgene expressing specifically a cytoplasmic GFP in muscles (marker HBR4). Genotypes are indicated above each series. Muscles detach from the outer body wall in *vab-10*(Δ*SH3*); *pak-1*(*RNAi*) embryos (arrows). Scale bar, 10 μm.

**Figure S3.**
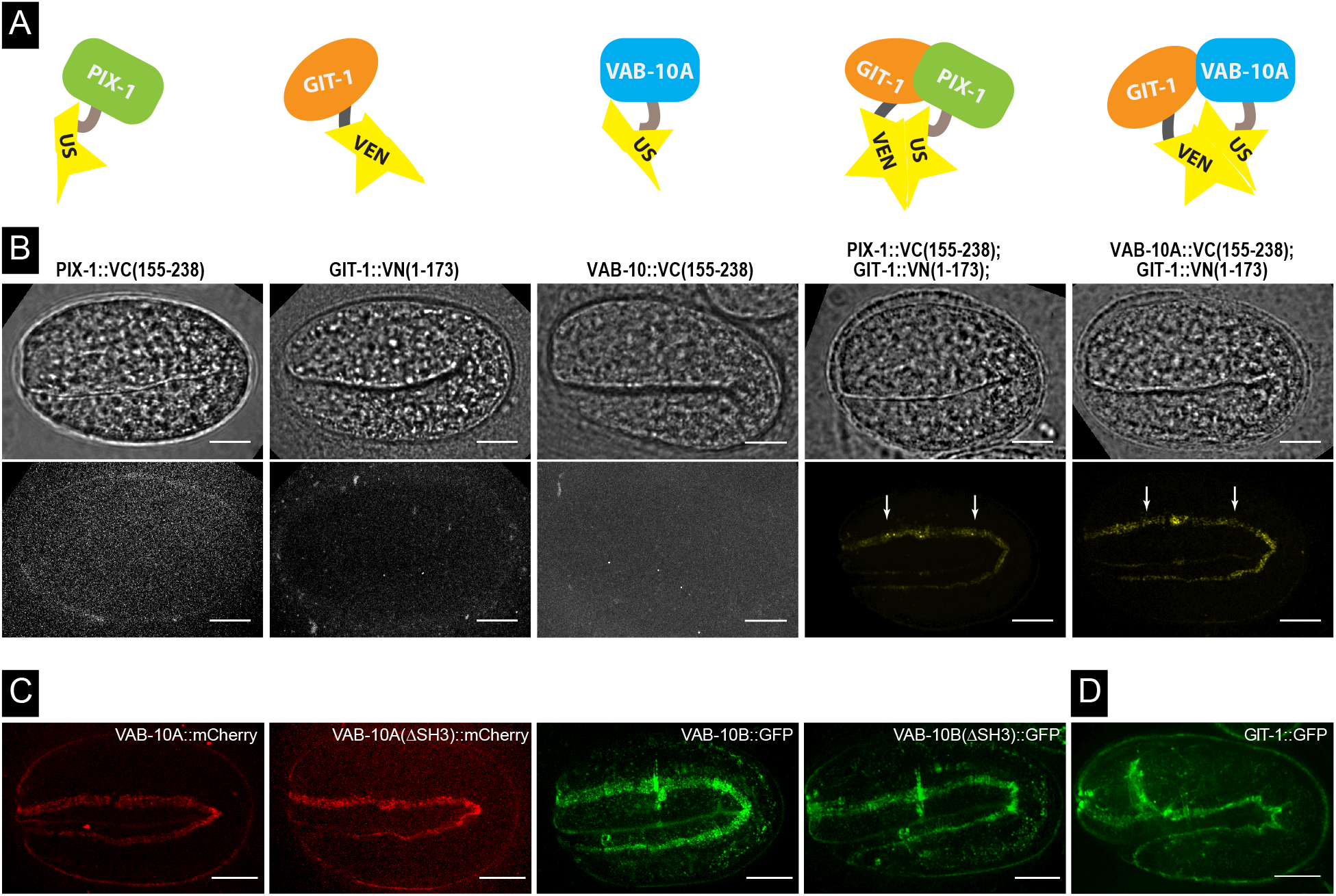
VAB-10A and GIT-1 localize within less than 10 nm distance from each other. **(A)** Strategy for the bi-fluorescence complementation strategy to define whether VAB-10A and GIT-1 are in close proximity. VEN, fusion with residues 1-173 of the Venus protein; US, fusion with residues 155-238 of the Venus protein. **(B)** Wide-field (upper panels) and spinning disc fluorescence (lower panels) micrographs of embryos expressing PIX-1::Venus(155-238), GIT-1::Venus(1-173), VAB-10A::Venus(155-238), GIT-1::Venus(1-173) with PIX-1::Venus(155-238), GIT-1::Venus(1-173) with VAB-10A::Venus(155-238). Note the clear hemidesmosomal pattern in the last two panels (arrows). **(C)** Spinning disc fluorescence images of 2-fold embryos homozygous for *vab-10A*::*mCherry* (*mc100*), *vab-10A*(Δ*SH3*)::*mCherry* (*mc109*), *vab-10B*::*GFP* (*mc123*), or *vab-10B*(Δ*SH3*)::*GFP* (*mc124*). The pattern of VAB-10A and VAB-10B was identical to that observed with antibodies (Bosher et al., 2003), and remained unaffected by the Δ*SH3* deletion. (**D**) Spinning disc fluorescence images of a 2-fold embryo homozygous for the CRISPR-generated knockin *git-1*::*GFP* (*mc86*). Scale bars, 10 μm.

**Figure S4.**
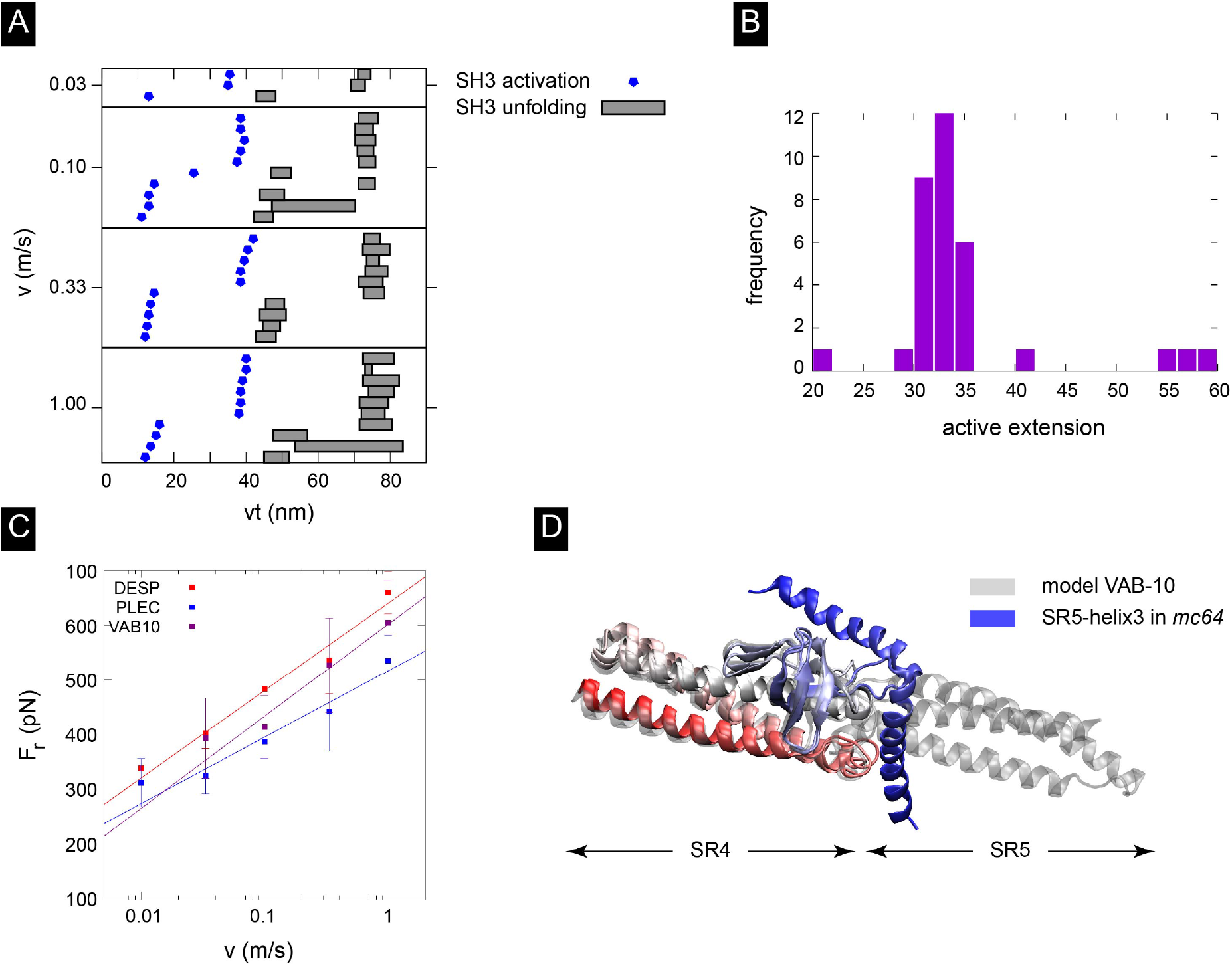
Quantification of the molecular dynamics simulations for VAB-10A plakin domain. Molecular dynamics simulation data on which Fig. 6 is based. **(A)** Points of SH3 activation and SH3 unfolding as a function of total extension (v†), depending on the pulling velocity. In all simulations, the activation and unfolding are decoupled events and activation happens first. **(B)** The frequency of the active extension of VAB10. For example, 30 nm means that after SH3 domain activation, another 30 nm of extension is required for the SH3 domain to start unfolding. **(C)** Comparison of rupture forces between plectin, desmoplakin and VAB-10. Note that VAB-10 appears to show an intermediate rupture force between plectin and desmoplakin. **(D)** Predicted 3D structure of the SR4-SR(+SH3)5 domain of VAB-10 (grey), and that of the same area for the mutant VAB-10(mc64) protein lacking the first two helices of SR5. The prediction is that the SH3 domain could still bind to SR4, but that most likely the remaining helix3 of SR5 would be destabilized (shown in two orientations).

## Captions for movies

**Movie S1**: DIC time-lapse video of a *pak-1*(*RNAi*) embryo during its active elongation.

**Movie S2**: DIC time-lapse video of a *vab-10*(Δ*SH3*); *pak-1*(*RNAi*) embryo during elongation.

**Movie S3**: DIC time-lapse video of a *vab-10*(Δ*SH3*); *git-1*(*tm1962*) embryo during elongation.

**Movie S4**: Fluorescence time-lapse video of a *pak-1*(*RNAi*) embryo homozygous for *let-805*::*gfp* during elongation.

**Movie S5**: Fluorescence time-lapse video of a *vab-10*(Δ*SH3*); *pak-1*(*RNAi*) embryo homozygous for *let-805*::*gfp* during elongation.

**Movie S6**: Fluorescence time-lapse video of a *vab-10*(Δ*SH3*)::*mCherry*; *pak-1*(*RNAi*) embryo homozygous for *let-805*::*gfp* during elongation.

**Movie S7**: Combined fluorescence video of embryos homozygous for *git-1*::*gfp* after knocking down the genes indicated on the movie.

Movies S1-S3 correspond to a single focal plane, Movies S4-S7 to the full projection. The frame rate is 30 frames/second (Movies S1-S3), 15 frames/second (Movies S4-S6) and 6 frames/second (Movie S7).

---

**Table S1.**
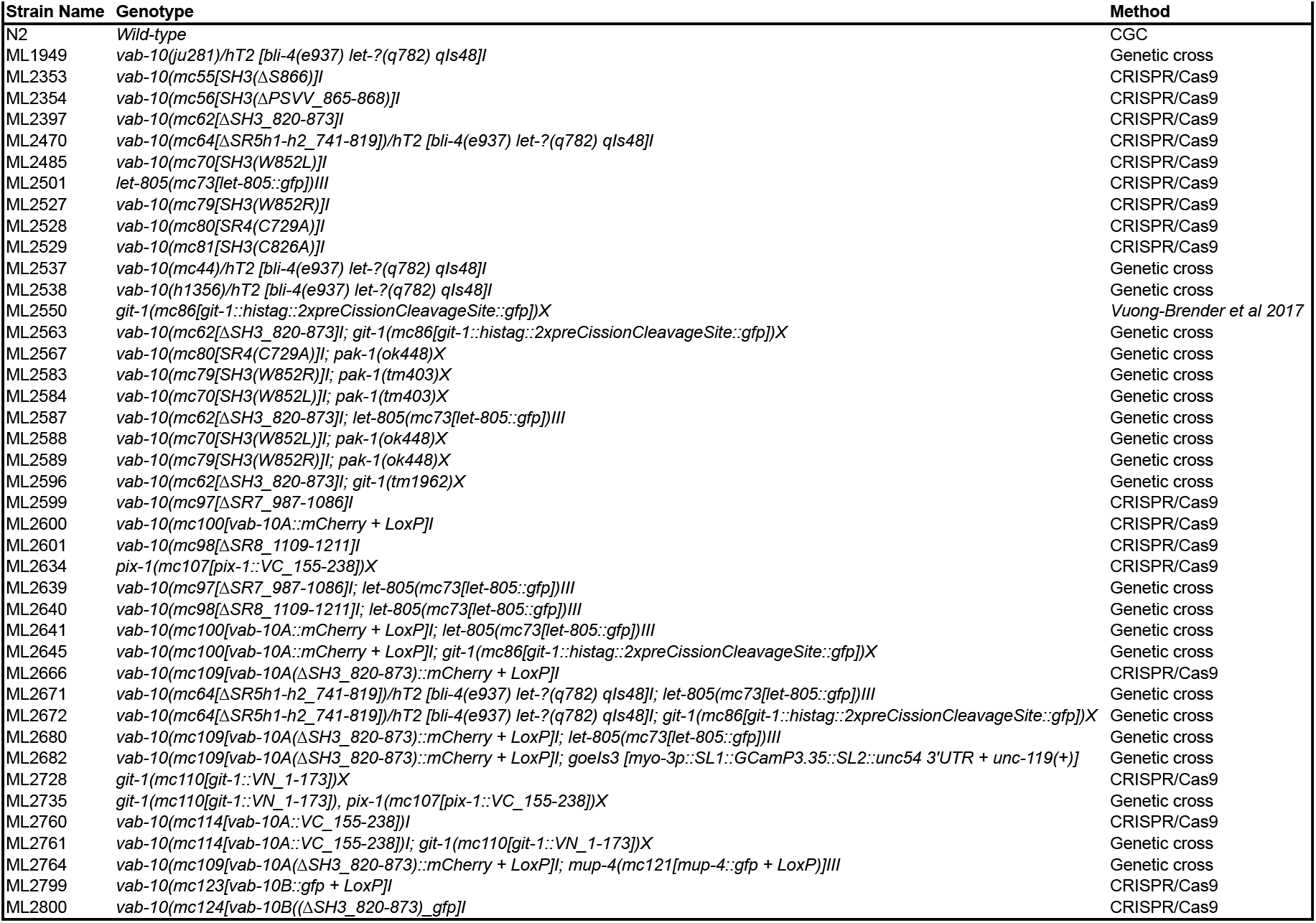

